# A general role for GGA adaptors in the modulation of AP-1-dependent trafficking

**DOI:** 10.64898/2026.03.25.714221

**Authors:** Alexander Stockhammer, Antonia Klemt, Aline Daberkow, Jelena Mijatovic, Laila S. Benz, Christian Freund, Benno Kuropka, Francesca Bottanelli

**Affiliations:** Institute of Chemistry and Biochemistry, Department of Membrane Trafficking, Freie Universität Berlin, Thielallee 63, 14195 Berlin, Germany; Osnabrück University, Department of Biology/Chemistry, Biochemistry section, Barbarastrasse 13, 49076 Osnabrück, Germany; Institute of Chemistry and Biochemistry, Department of Protein Biochemistry, Freie Universität Berlin, Thielallee 63, 14195 Berlin, Germany

## Abstract

The Golgi-localized, γ-ear containing, ADP-ribosylation factor binding proteins (GGAs) are a family of adaptor proteins that regulate transport of specific cargo receptors from the Golgi to endosomes. For many years it was assumed that GGAs transport cargo via interaction with the adaptor complex AP-1. However, recent findings suggest that GGA and AP-1 may have opposing roles, with GGAs facilitating forward transport between Golgi and endosomes, and AP-1 mediating the opposite trafficking step. To shed light on the functional connection of GGAs with AP-1, we combined CRISPR-Cas9 gene editing with live-cell imaging and TurboID-based proximity labelling. We find that GGAs localize not only to the Golgi apparatus but also, to a greater extent, to peripheral ARF1-positive compartments responsible for secretory trafficking and endocytic recycling. At both, the Golgi and peripheral sites, we observe distinct sorting domains containing either AP-1 or GGAs alone, as well as domains in which both adaptors are present. Interestingly, GGAs can recruit clathrin lattices independently of AP-1. Proximome mapping shows that AP-1 specific cargoes only localize to AP-1 domains in the absence of GGAs. These findings point to a regulatory role of GGAs in AP-1 transport. We speculate that GGAs prevent binding of AP-1 to its cargo clients to avoid premature retrieval and to modulate bi-directional trafficking between the Golgi and endosomes.

## Introduction

In eukaryotic cells, a complex protein machinery directs cargo flow between the Golgi, trans-Golgi-network (TGN), different classes of endosomes and the plasma membrane^1^. Adaptor proteins facilitate selection, enrichment, and transport of various secretory and endocytic cargoes via interacting with the cytoplasmic tails of receptors and transmembrane cargoes^2^. The small GTPase ADP-ribosylation factor 1 (ARF1) recruits the adaptor protein complexes AP-1, AP-3 and AP-4 as well as the group of monomeric Golgi-localized, γ-ear containing, ADP-ribosylation factor binding proteins (GGAs) to the TGN and endosomal membranes^2–4^. Current evidence suggests that GGAs mediate the transport of cargo receptors, such as the mannose-6-phosphate receptor (M6PR) and sortilin, from the trans-Golgi network (TGN) to endosomes^5–7^.

Vertebrates express three different GGA proteins (GGA1, GGA2 and GGA3 in human). GGA1-3 share a similar domain structure, yet they differ in their ubiquitin-binding properties and mechanisms of auto-inhibition^8^. It remains unclear whether they function redundantly or possess distinct specialized sorting roles. Knock out (KO) data from mice showing that absence from GGA2, or the combined loss of GGA1 and GGA3, led to increased neonatal lethality^9^, suggest that at least GGA2 plays a non-redundant role in intracellular transport. A triple GGA KO impairs the transport of hydrolases from the Golgi to endosomes and traps their receptor M6PR at the TGN^10^ and mutations in the GGA-sorting signal of the M6PR cause its retention in the Golgi^11^. These findings support a role for GGAs in transport of hydrolases to endo-lysosomal compartment via their receptor M6PR. In contrast, loss of AP-1 results in trapping of the M6PR on endosomal membranes^12,13^ indicating that AP-1 is required for the retrieval of M6PR to the Golgi. This has led to a model in which GGAs mediate the anterograde transport of hydrolases, while AP-1 facilitates the retrograde retrieval of their receptors back to the TGN^14^. However, GGAs were shown to interact with AP-1^15^, and while such a model would account for the observed trafficking defects, the functional necessity of an AP-1/GGA interaction remains unclear. Because both AP-1 and GGAs bind the coat protein clathrin^16,17^, it has been suggested that both adaptors jointly facilitate the formation of clathrin-coated vesicle (CCV) for intracellular transport. Importantly, GGA2 was found to be strongly depleted from CCVs in AP-1 knock-sideway experiments, suggesting that the interaction with AP-1 is needed for its packing into clathrin coats^18^. In yeast, AP-1 and GGAs are recruited to the membrane independently^19^ and it has been proposed that GGAs recruitment precedes AP-1 association to the membrane^20,21^, supporting a model in which the two adaptors act sequentially. These diverse observations have led to distinct models proposing how the two adaptors function in protein transport between the TGN and endosomes^8,14^. However, none of the models can fully explain the observed phenotypes nor can they explain the opposed directionality in transport that was proposed for GGAs and AP-1^14,18^.

In this study, we endogenously tag the three mammalian GGAs to investigate their localization and functional relationship with AP-1 in living cells. We focus on the spatial and functional interplay of GGAs and AP-1, in relationship to ARF1-positive tubular vesicular carriers which we have recently shown to drive biosynthetic and endocytic recycling cargo flux^22^. Finally, we utilize comparative TurboID-based proximome mapping to reveal differences and similarities in GGAs and AP-1 nanoscale environment, to ultimately understand their role in intracellular transport. Our results suggests that GGAs play a general regulatory role in AP-1 mediated protein transport.

## Results

### All three GGAs are recruited to the same nanodomains on the TGN and peripheral ARF1-positive compartments

GGAs associate to membranes via interaction with the small GTPase ARF1^4^. Recently, we have found that different classes of ARF1-positive tubular-vesicular compartments direct secretory trafficking from the TGN to the plasma membrane as well as endocytic recycling in the cellular periphery^22^. Although GGAs were primarily detected at Golgi and TGN, they were also reported to populate endosomal membranes including recycling endosomes^23–26^. So far, all studies to assess GGA localization were done using either fluorescently labelled fusion proteins that were transiently expressed in cells or by staining the endogenous proteins using antibodies in fixed cells. However, both overexpression and fixation can cause artifacts and interfere with protein localization^27–29^. Thus, to get a better understanding of the role of GGAs in protein transport, we first set out to reveal the intracellular localization of endogenous GGAs in living cells.

To test GGAs localization in unfixed and unperturbed cells, we used the CRISPR-Cas9 system to endogenously tag GGA1-3 with eGFP in HeLa cells expressing ARF1^EN^-Halo (Fig. 1). We found that all three GGAs localized to the Golgi apparatus as reported in the literature (Supplementary Fig. 1). Interestingly, GGAs also localized to peripheral ARF1-positive compartments (Fig. 1A). Their localization to ARF1 compartments becomes evident when focusing on a plane near the basal membrane of the cell (Fig. 1B). When examining the cellular plane that most clearly resolves the Golgi and TGN, GGAs are observed to localize predominantly to these membranes, while peripheral localization is comparatively infrequent (Fig. 1B). Quantification of signal intensity of single eGFP-GGA2 domains on the Golgi versus GGA2 domain on periphery compartments, showed that GGA2 domains at the Golgi are ∼ 2-fold brighter compared to those in the periphery (Fig. 1C). The presence of GGA domains on peripheral ARF1 compartments indicates that GGAs may have a broader role in trafficking beyond TGN export alone.

**Figure 1:**
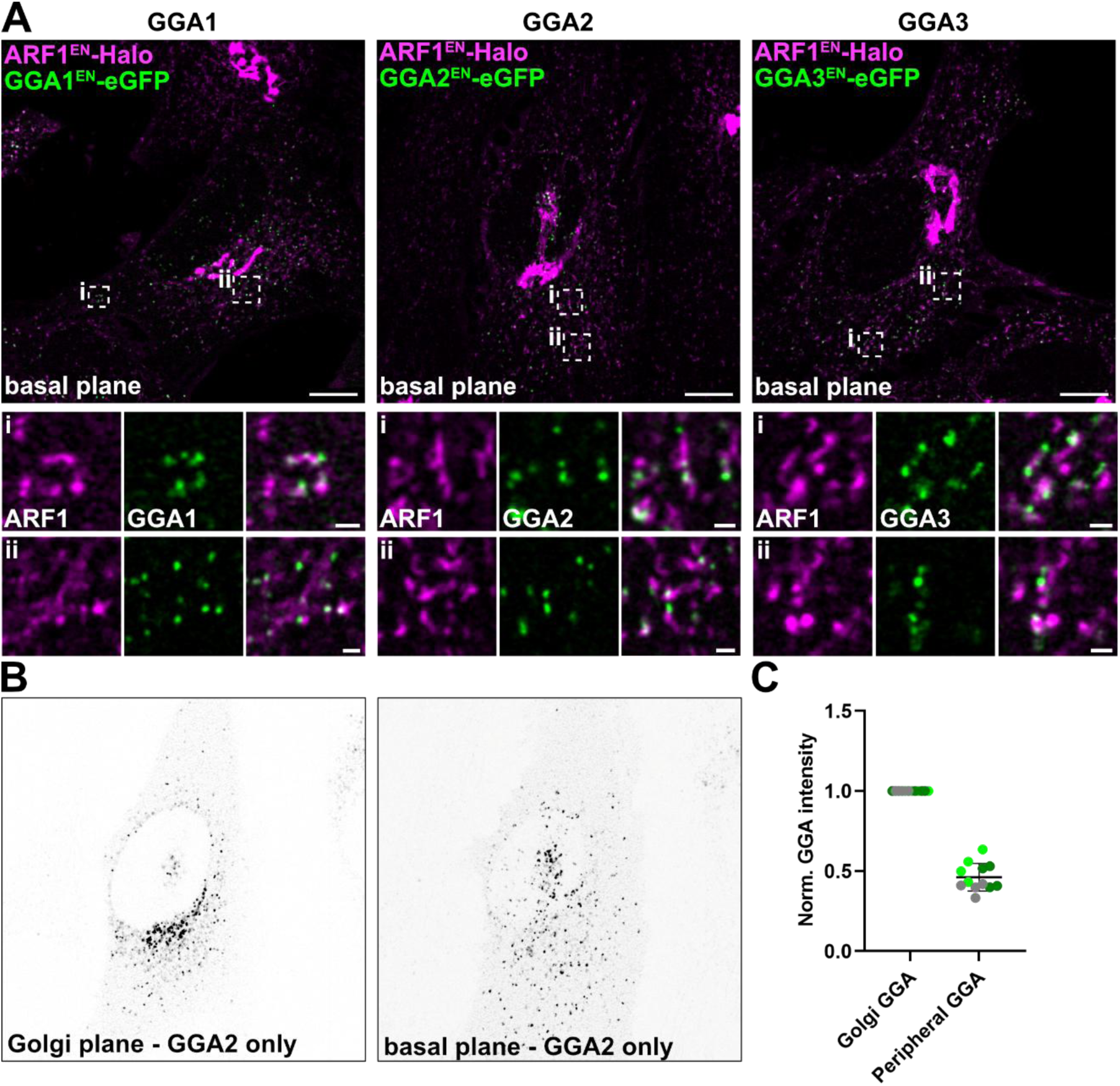
GGAs localize to the Golgi and peripheral ARF1 compartments. (**A**) Live-cell confocal imaging of GGA1^EN^-eGFP/ARF1^EN^-Halo, GGA2^EN^-eGFP/ARF1^EN^-Halo and GGA3^EN^-eGFP/ARF1^EN^-Halo HeLa cells labeled with CA-JFX_650_ shows GGAs association to peripheral ARF1 compartments. (**B**) Using GGA2^EN^-eGFP/ARF1^EN^-Halo HeLa cells as example, GGA2 distribution at different confocal focal planes is shown, highlighting that at the Golgi plane GGA2 localizes primarily to the Golgi (ARF1 signal from **A** can be used as reference for Golgi-localization) while at a basal plane GGA2 is dispersed throughout the cellular periphery. (**C**) Quantification of signal intensity of GGA2 punctae at Golgi and ARF1 compartments in GGA2^EN^-eGFP/ARF1^EN^-Halo HeLa cells. In total, the signal intensities from 10 GGA punctae per category were averaged in 12 cells from 3 independent experiments and normalized to the averaged signal intensities of TGN associated GGA punctae. Replicates are shown in different colors and each dot represents a single cell of the replicate. CA=chloroalkane (HaloTag substrate). Scale bars: 10 μm (overview) and 1 μm (crops).

Next, with our gene editing tools in hand we wanted to test the extent of co-localization of the three GGAs isoforms. In fixed cells, GGAs largely colocalize to the Golgi^30^, but immunofluorescence largely missed the pool of GGAs localizing to peripheral ARF1 compartments. Therefore, we created double knock in (KI) cell lines with different combinations of GGAs tagged with either HaloTag or eGFP (Fig. 2) for colocalization analysis. We observed perfect colocalization of different GGAs in domains either in the Golgi area (Fig. 2A-Ci) and in the cellular periphery (Fig. 2A-Cii). Based on these data, we conclude that all GGAs are recruited to shared nanodomains at the Golgi and in the cell periphery, where they coordinate transport.

**Figure 2:**
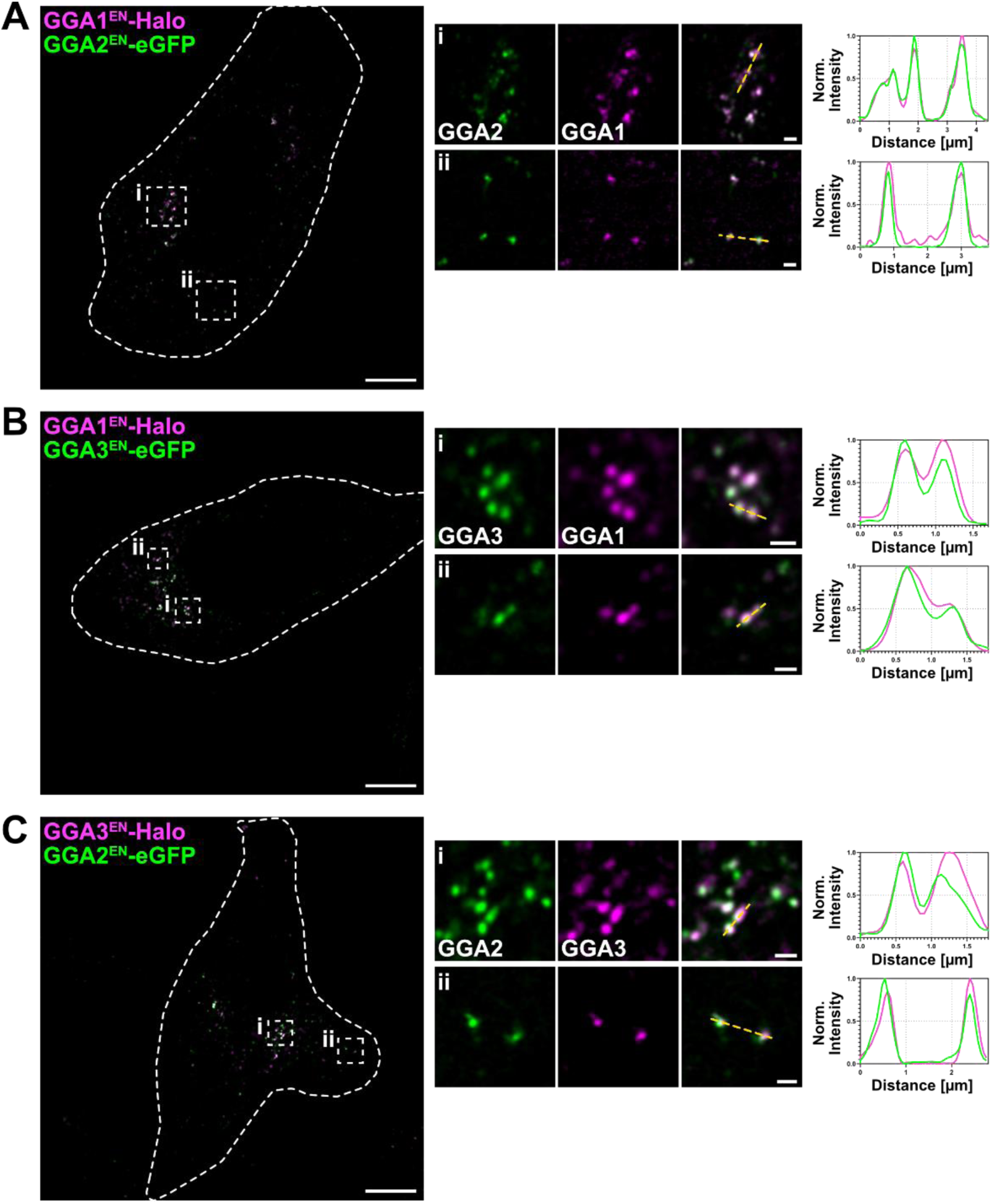
All GGA protein localize to the same sorting domains. (**A-C**) Live-cell confocal imaging of GGA2^EN^-eGFP/GGA1^EN^-Halo (**A**), GGA3^EN^-eGFP/GGA1^EN^-Halo (**B**) and GGA2^EN^-eGFP/GGA3^EN^-Halo (**C**) HeLa cells labeled with CA-JFX_650_ highlights colocalization of GGAs in the in the perinuclear area and in the cellular periphery. Examples of different compartments and line profiles showing perfect colocalization of the different GGAs. Scale bars: 10 μm (overview) and 1 μm (crops).

### AP-1 and GGAs localize to distinct and shared membrane domains

AP-1 was shown to interact with GGAs^15^ and to localize to the Golgi and peripheral ARF1 compartments^22^ similarly to what we have observed for GGAs (Fig. 1-2). As the functional and physical association of these adaptors remains unclear, we next sought to define their membrane association relative to one another on Golgi and peripheral ARF1-positive compartments.

To address this, we generated a triple KI cell line enabling the simultaneous visualization of AP-1 and GGA2 alongside the underlying ARF1-positive membranes (HeLa ARF1^EN^-eGFP/AP1µA^EN^-SNAP/GGA2^EN^-Halo) (Fig. 3A). Intriguingly, we found three distinct populations of nanodomains with different adaptors composition on Golgi and on peripheral ARF1 compartments (Fig. 3B-C). We observed domains containing either AP-1 or GGA2, (Fig. 3B-C highlighted with white arrows) as well as domains containing both adaptors (Fig. 3B-C highlighted with red arrows). This suggests the existence of functionally distinct AP-1 and GGA nanodomains in addition to AP-1/GGA domains. Moreover, dynamic live-cell imaging highlighted AP-1 dissociating from AP-1/GGA-positive nanodomains or being recruited to a GGA2 nanodomain (Fig. 3D-E). Conversely, we could also observe GGA2 dissociating from an AP-1/GGA2-positive domain (Fig. 3E), suggesting dynamic exchange of adaptors on pre-assembled membrane domains of both kinds (AP-1 and GGA2). The same experiments were also carried out in cell lines expressing endogenously tagged GGA1 and GGA3 which displaying the same localization and dynamics (Supplementary Fig. 2). Subsequent dynamic imaging experiments were thus carried out with the GGA2 KI cell line.

**Figure 3:**
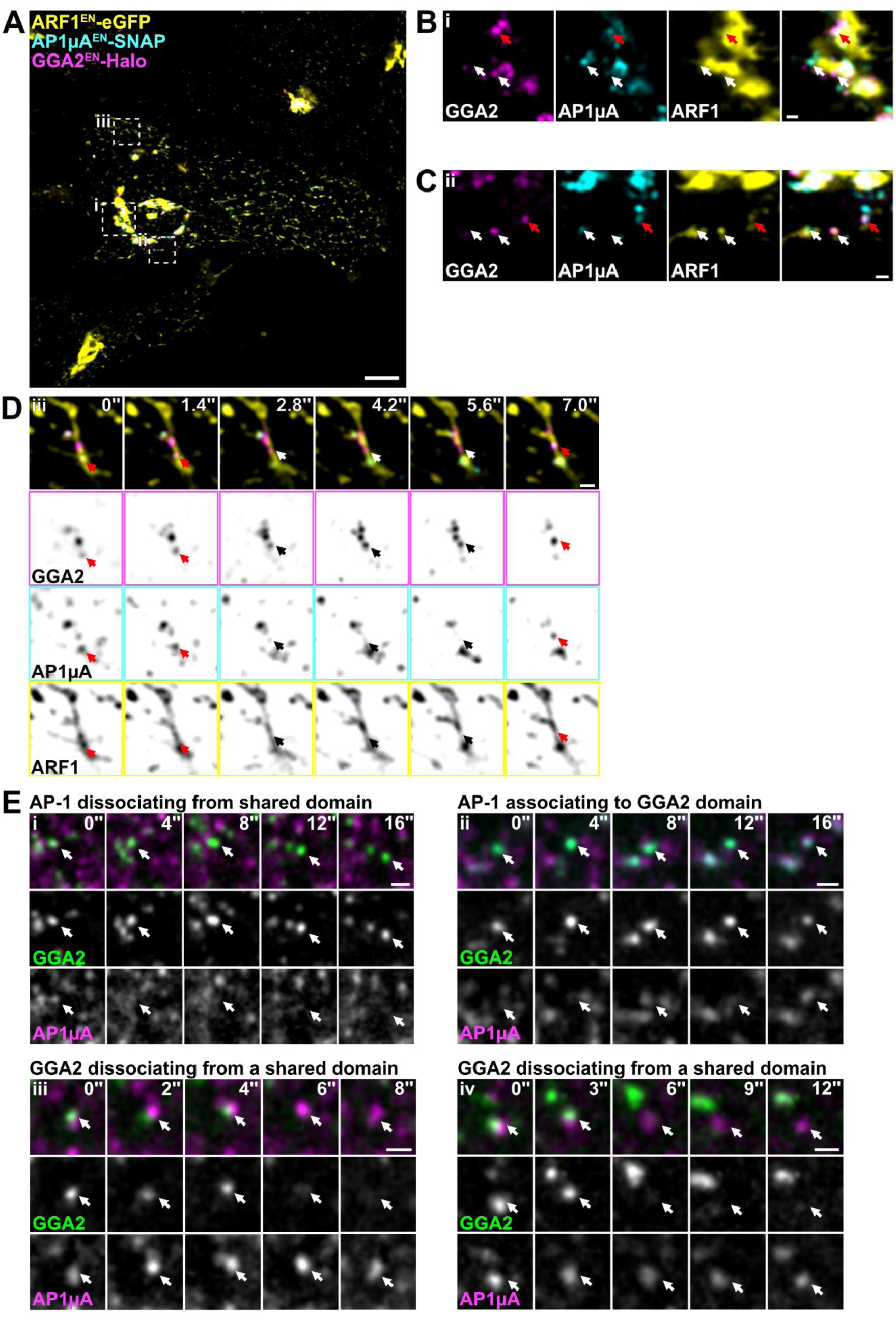
Transient association and dissociation of AP-1 and GGA from shared membrane domains. (**A**) Live-cell spinning disk imaging of ARF1^EN^-eGFP/AP1µA^EN^-SNAP/GGA2^EN^-Halo HeLa cells labeled with BG-JF_552_ and CA-JFX_650_. (**B-C**) Magnifications highlight domains at the TGN and at peripheral ARF1 compartments where only one of the two adaptors AP-1 or GGA2 (highlighted with white arrows) is present. Also, domains were both adaptors were present are shown (highlighted with red arrows). (**D**) Timelapse imaging revealed domain identity switches. Specifically, over time AP-1 is depleted from shared domains (red arrow in frames 0’’ – 2.8’’) or was found to associate to a GGA2 domain (red arrow in frame 7.0’’). (**E**) Additional timelapse imaging examples of GGA2^EN^-mStayGold/AP1µA^EN^-SNAP HeLa cells labeled with BG-JF_552_ showing selected frames of events where AP-1 associates and dissociates (i-ii) from GGA2 sorting domains and vice versa examples of GGA2 dissociating from AP-1 domains (iii-iv). BG=benzylguanine (SNAP-tag substrate). Scale bars: 10 μm (overview) and 1 μm (crops).

### AP-1 and GGAs colocalize on secretory and endocytic compartments

The most recent model proposes that AP-1 and GGAs perform opposing roles in anterograde transport from the TGN and retrograde transport, respectively^14^. According to such model, GGAs would predominantly localize to secretory TGN-derived carriers. As AP-1 domains were observed on TGN-derived ARF1 compartments involved in secretory trafficking and ARF1 compartments that are involved in endocytic recycling^22^, we wondered if the formation of GGA-AP1 sorting domains is limited to the TGN and anterograde TGN-derived carriers or also occurs on endocytic recycling compartments. First, to be able to visualize adaptor recruitment on TGN-derived secretory carriers we employed the retention using selective hooks (RUSH)^31^ system in combination with a M6PR-RUSH reporter, a known GGA client (Fig. 4A)^11,32^. The M6PR reporter is retained in the endoplasmic reticulum (ER) and is released upon the addition of biotin. M6PR is seen exiting the Golgi in vesicular carriers as previously reported^11,14^. In addition, we could observe M6PR-positive tubular carriers (Fig. 4A) decorated with AP-1 and GGA2, which localized mostly to shared sorting domains (arrows in Fig.4Ai).

**Figure 4:**
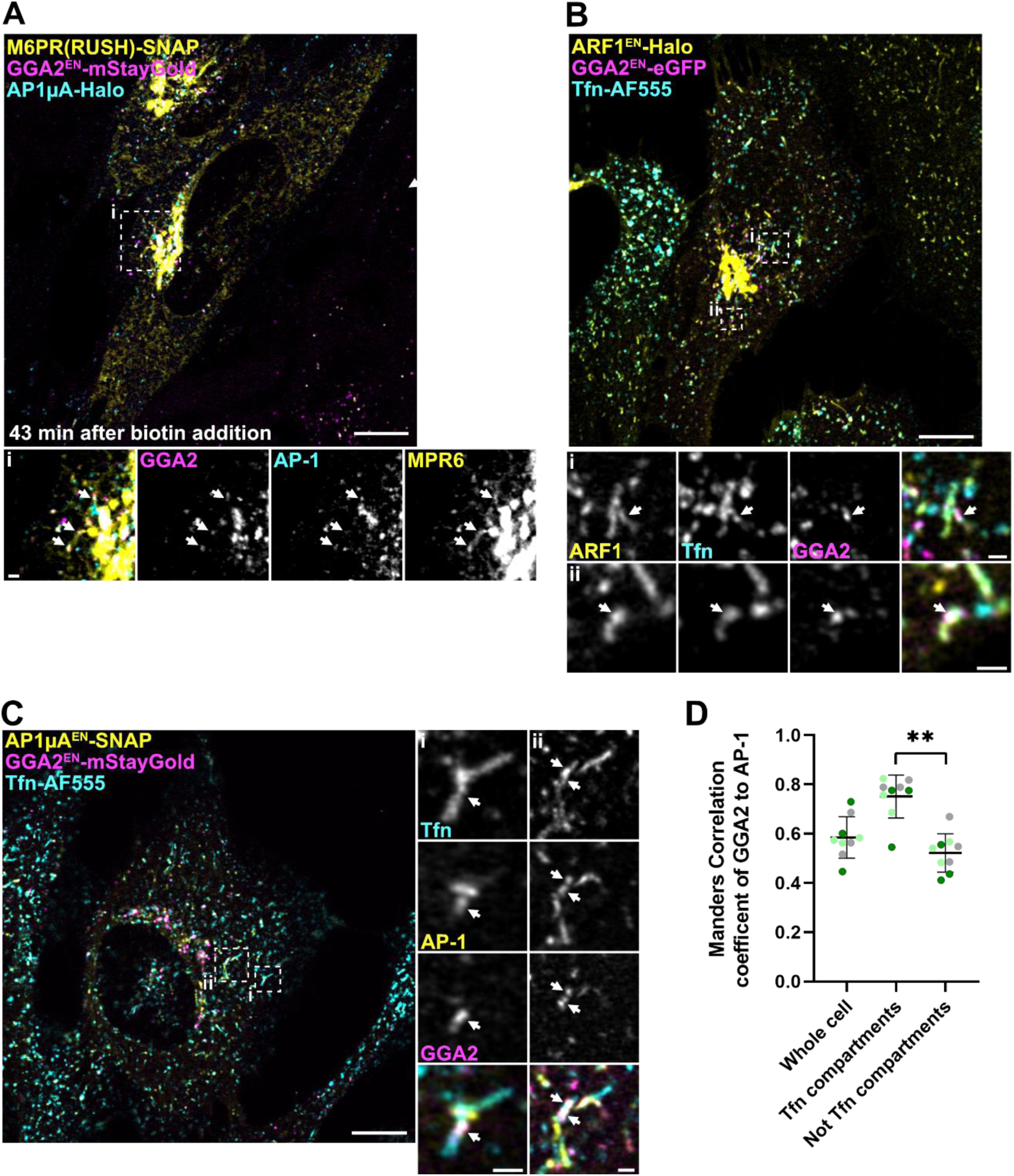
GGAs are recruited to secretory and recycling compartments. (**A**) GGA2^EN^-mStayGold/AP1µA^EN^-Halo HeLa cells transiently expressing the RUSH constructs streptavidin-SBP-SNAP-M6PR (M6PR(RUSH)-SNAP) labelled with CA-JF_552_ and BG-JFX_650_ were imaged with confocal microscopy after addition of biotin. A qualitative example shows the M6PR(RUSH) in TGN-derived tubules. Both adaptors GGA2 and AP-1 colocalize to specific domains on the TGN-derived tubules (highlighted with arrows) (**B**) Transferrin (Tfn) recycling assays were performed using fluorescently labelled Tfn (Tfn-AlexaFluor555). Live-cell confocal imaging of GGA2^EN^-eGFP/ARF1^EN^-Halo HeLa cells with CA-JFX_650_ shows GGA2 on ARF1 compartments which are filled with Tfn 5 min after addition of Tfn. (**C**) Live-cell confocal microscopy of Tfn recycling using Tfn-AlexaFluor555 in GGA2^EN^-mStayGold/AP1µA^EN^-SNAP HeLa cells labelled with BG-JFX_650_. The two adaptors AP-1 and GGA2 were found on Tfn-filled compartments. (**D**) Colocalization analysis using the Manders correlation coefficient reveals that colocalization of GGA2 and AP-1 is significantly increased on Tfn-filled compartments compared to Tfn negative membranes. In total 9 cells from three independent experiments were analysed, replicates are shown in different colours each dot representing a single cell. *P* value of two-sided *t*-test from averages of the biological replicates is <0.0051. Scale bars: 10 μm (overviews in) and 1 μm (crops).

To investigate the localization of GGAs on endocytic recycling compartments we performed transferrin (Tfn) uptake experiments using fluorescently-labeled Tfn. After internalization, Tfn was found in ARF1 compartments which harboured GGA2 domains (Fig. 4B). When we performed the same experiments in HeLa AP1µA ^EN^-SNAP/GGA2^EN^-mStayGold cells, we observed that GGA2 frequently colocalizes with AP-1 on Tfn-positive compartment (Fig. 4C, arrows highlight domains where GGA2 and AP-1 are present). These experiments show that GGAs not only control cargo export from the Golgi, but also act on endosomes involved in endocytic recycling. To understand whether GGA2/AP-1 association may be different in the context of regulation of TGN export and endocytic recycling, we quantified the amount of GGA colocalizing with AP-1 domains on both secretory (Tfn-negative) and endocytic (Tfn-positive) compartments. Interestingly, the colocalization of GGA2 and AP-1 significantly increases on Tfn-filled compartments compared to compartments which are not involved in endocytic recycling (Fig. 4D). This suggests that the association between GGA2 and AP-1 becomes stronger on membranes involved in endocytic recycling.

### GGAs form clathrin lattices independent from AP-1

The presence of heterogeneous nanodomains with varying adaptor compositions prompted us to investigate their ability to recruit the clathrin coat. GGAs bind to clathrin via their hinge and ear domains^17^ and *in vitro* assays showed that GGAs were sufficient to recruit clathrin to liposomes^33^ . However, GGAs bind clathrin more weakly than AP-1^17^ and knock-sideways experiments of AP-1 led to strong depletion of GGA2 from CCVs^18^, suggesting that AP-1 might be required for clathrin lattice formation in cells. As we observed association and dissociation of GGAs from AP-1 sorting domains (Fig. 3), we were wondering whether GGAs bind clathrin in the absence of AP-1.

We thus created a HeLa GGA2^EN^-mStayGold/AP1µA^EN^-SNAP/Halo-CLCa^EN^ cell line to be able to capture the spatial arrangement of GGA2 in comparison to AP-1 and clathrin in living cells (Fig. 5). As expected, clathrin was associated with the entirety of AP-1 nanodomains. Intriguingly, we could detect GGA2 domains devoid of AP-1 that are either positive or devoid of clathrin (Fig. 5A). Overall, we identified three different populations of GGA domains; (1) GGA domains that are devoid of AP-1 and clathrin (Fig. 5A, highlighted with red arrows), (2) GGA domains positive for clathrin but devoid of AP-1 (Fig. 5A, highlighted with white arrows), (3) GGA domains positive for both clathrin and AP-1 (Fig. 5A, highlighted with yellow arrows). Quantification shows that the majority of GGA domains (∼70%) also harbour AP-1 and clathrin and only a small percentile of GGA domains is devoid of both AP-1 and clathrin (Fig. 5B). These data suggest, that GGAs are able to form clathrin lattices independently from AP-1 *in vivo*. However, the fact that we observed GGA2 domains devoid of AP-1 and clathrin indicates that the clathrin lattice is not required to assemble GGAs on membranes. It is possible that these domains are newly formed sorting domains that will acquire clathrin over time. Moreover, our data shows that the majority of GGAs co-localizes with AP-1, supporting a collaborative role of both adaptors in protein transport.

**Figure 5:**
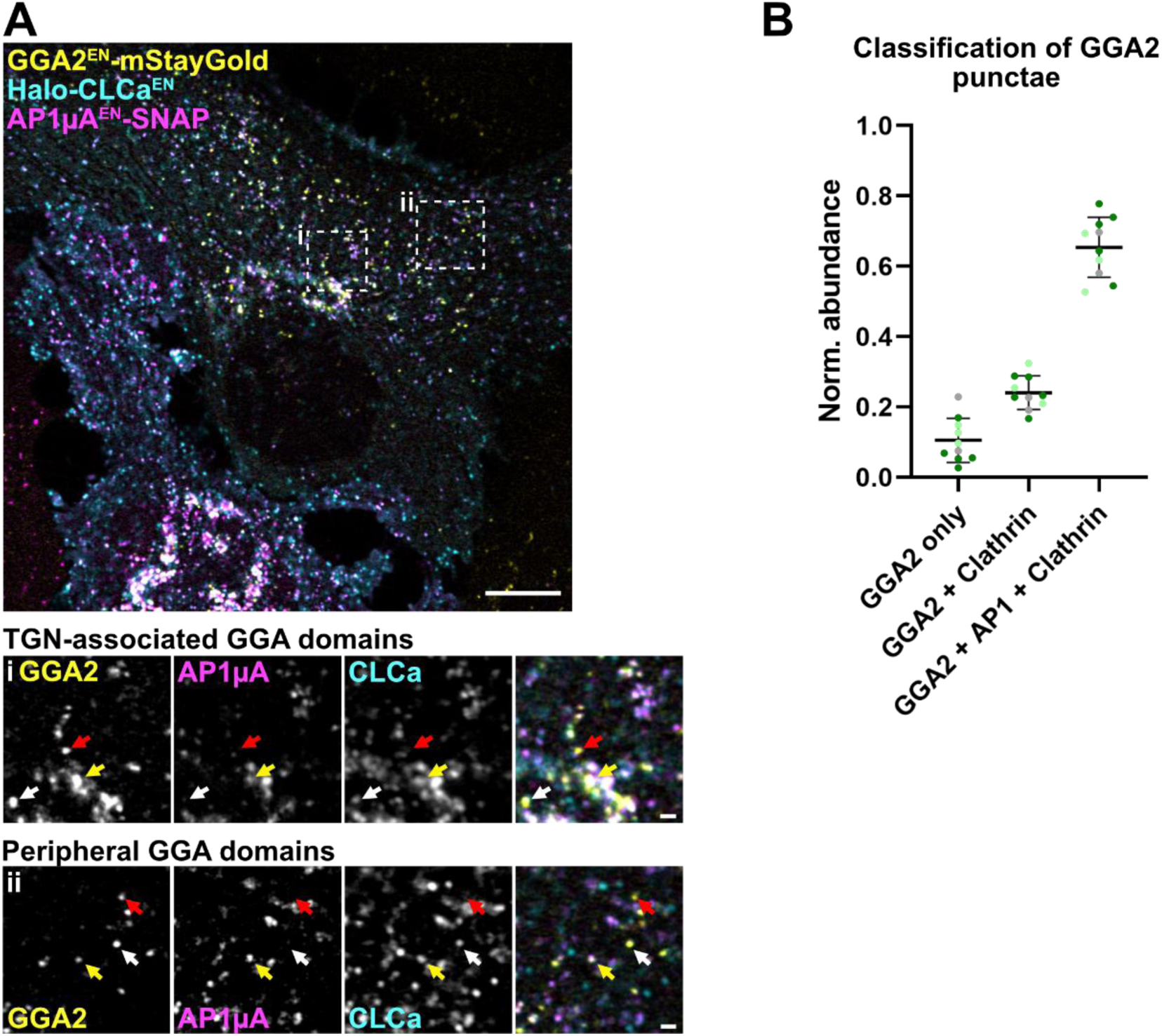
GGAs can bind clathrin independent of AP-1. (**A**) Live-cell confocal imaging of GGA2^EN^-mStayGold/AP1µA^EN^-SNAP/Halo-CLCa^EN^ HeLa cells labeled with BG-JF_552_ and CA-JFX_650_ reveals three distinct GGA2-populations based on its association with clathrin and AP-1. Examples for the three populations which are found at the TGN and in the periphery of the cell are highlighted by arrows. GGA2 that is neither associated with AP-1 nor clathrin is highlighted with red arrows, GGA2 that binds clathrin but not AP-1 is highlighted with white arrows and GGA2 found associated with both clathrin and AP-1 is highlighted by yellow arrows. (**B**) Quantification of the different GGA2 populations within in the cell. In total 10 cells were analysed; replicates are shown in different colours each dot representing a single cell. Scale bars: 10 μm (overview) and 1 μm (crops).

### Comparative proximome analysis reveals that AP-1-dependent cargoes are excluded from GGA nanodomains

Motivated by the observation that GGA domains can exist either with or without AP-1 (Fig. 2 and Fig. 4), we next sought to understand differences in molecular composition between AP-1-only, GGA-only and AP-1/GGA-domains. For this, we opted for a TurboID-based proximome mapping approach^34,35^ which allowed to identify proteins which are recruited specifically to domains where either only AP-1 or GGAs are present (Fig. 6). We endogenously tagged the three GGAs with TurboID and used an established HeLa AP1µA^EN^-TurboID cell line^35^ to obtain GGAs and AP1s proximomes. To map and compare the proximome of GGAs and AP-1, we analyzed streptavidin-purified biotinylated proteins by MS using label-free quantification from the different KI cell lines. To identify proteins specifically recruited to GGA or AP-1 domains we compared their relative intensity between datasets of different GGAs and AP-1 (Fig. 6A-C). Surprisingly, we found that AP-1 specific cargo proteins such as the transferrin receptor (TFRC)^36^, the copper transporter ATP7A^37^ and integrins (e.g. ITGB1)^38^ were highly enriched in the AP-1 proximome dataset but were not detected in any GGA dataset. This indicates that these cargoes are only presents in AP-1 domains devoid of GGAs. A full list of proteins that we found enriched when we compared AP-1 interactome and GGA interactomes is provided in supplementary table 1. We also looked for proteins enriched in both the AP-1 and GGA TurboID datasets when compared to a WT control sample, as we suspected those proteins could be found in domains where and when both adaptors are present (Supplementary table 2). Among those proteins we find several proteins which may always be present in AP-1 sorting domains such as epsinR, aftiphilin and γ-synergin^39,40^. The M6PR is also on the list of proteins enriched in both AP-1 and GGAs TurboID datasets, suggesting both adaptors are needed for its sorting. While we assume it may be present in GGA/AP-1 sorting domains, it may also be sorted via AP-1 or GGA-only sorting domains. Lastly, only a few proteins were specifically enriched in the GGAs proximome in comparison to AP-1 (Supplementary table 3), which include the known GGA interacting proteins rabaptin-5 (RABEP1) and p63 (CCDC91)^41,30^. These proteins may preferentially bind to GGAs in the absence of AP-1 or could possibly play a regulatory role in the AP-1/GGA association.

**Figure 6:**
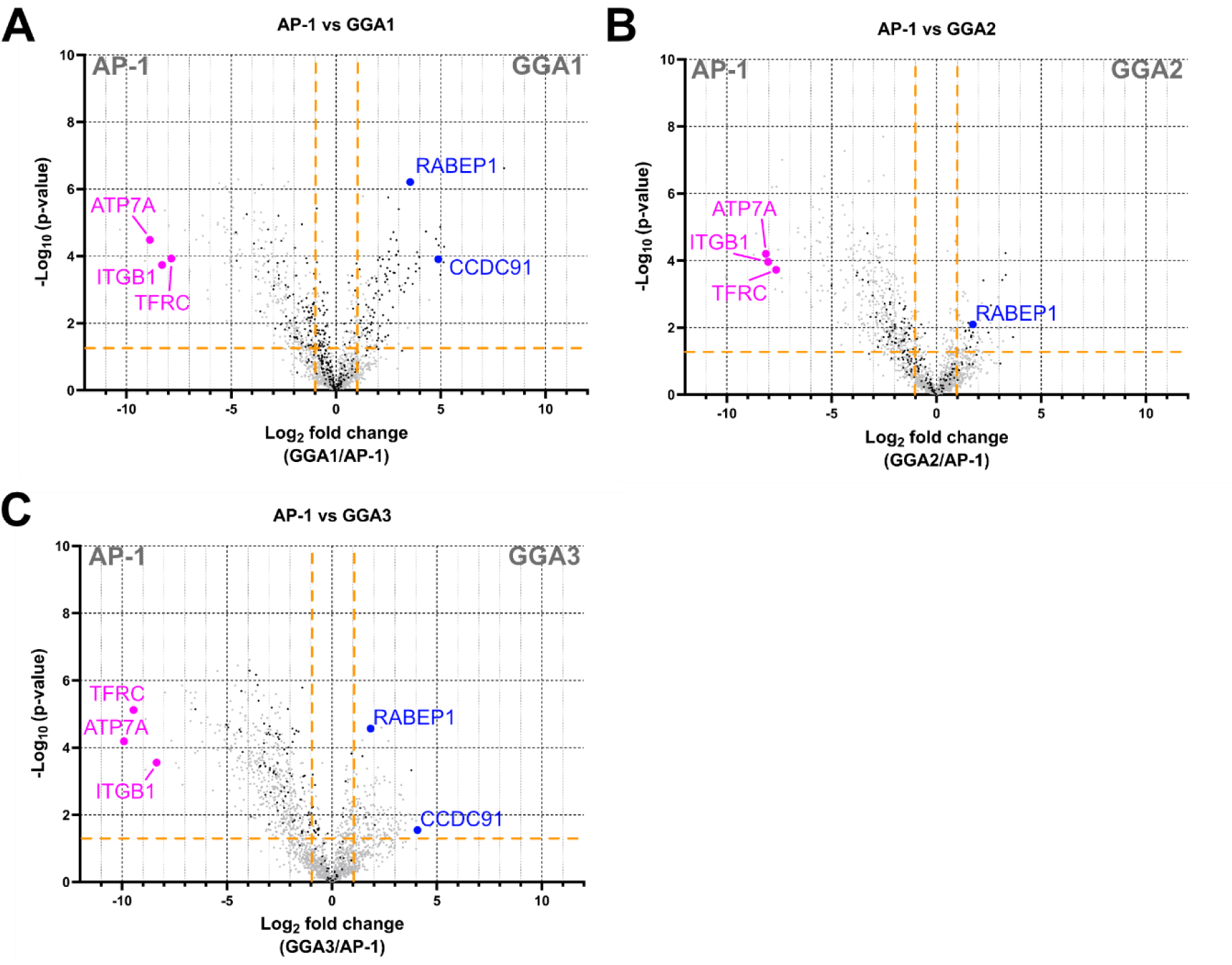
AP-1 only binds cargoes in the absence of GGA. (**A-C**) Volcano plot showing the changes in relative protein intensity between AP1µA^EN^–TurboID and GGA1^EN^–TurboID (A), GGA2^EN^–TurboID (B) and GGA3^EN^–TurboID HeLa cells. Proteins that show significant changes in their relative intensity are shown in the top left (AP-1) and top right (GGA) corner (*P*<0.05 and log_2_ fold change >1 or <−1) separated by the orange lines. Proteins that were not significantly enriched in GGA-TurboID samples compared to WT samples are shown in light grey. Selected AP-1-specific interactors are highlighted in magenta and selected GGA-specific interactors shown in blue.

Overall, our proximome analysis reveals that many AP-1 cargo proteins appear to only localize to AP-1 sorting domains in the absence of GGAs. On the contrary, we could not detect any cargo proteins that would localize exclusively to GGA-only sorting domains. While proximome mapping does not reveal any direct interactions, it allowed a comprehensive characterization of the composition of AP-1 or GGA-specific sorting domains.

## Discussion

Despite decades of investigation, it is still unclear how GGAs modulate intracellular transport. It is widely accepted that GGAs facilitate forward transport of hydrolases and their receptors from the TGN to endosomes^5,10,14,18^. Such directionality of transport would require primary localization of the GGAs to the TGN, where they were localized primarily in the past^30,32,42^, as coats are exclusively present on donor compartments where they assemble to select cargoes. However, a combination of endogenous tagging of GGAs and live-cell imaging enabled us to visualize a significant population of GGAs which localize to peripheral ARF1-positive compartments (Fig. 1). Importantly we were able to show that those ARF1 compartments are involved in both secretory and endocytic recycling (Fig. 4), suggesting that the role of GGAs in intracellular transport goes beyond protein export from the Golgi. The extent of GGAs localization to peripheral endosomes^30,43^ has been likely underestimated due to the fact that these ARF1 compartments are sensitive to fixation^44,28^ and many studies towards GGA localization were antibody-based^32,42,45^. Thus, the widespread localization of GGAs across various intracellular membranes suggests they play a broader role in trafficking.

But what would be function of GGAs on recycling compartments? Our data confirms the functional connection between GGA and AP-1 as we frequently observed domains with both adaptors present (Fig. 3). GGAs were visualized in the proximity of AP-1^18,35^ and the two adaptors were shown to physically interact^15^, supporting the idea that both adaptors would cooperate in transport^18,20^. However, while GGAs are important for export of specific proteins from the Golgi such as the MP6R^11^, AP-1 has been recently shown to be required for the retrieval of such proteins from endosomes to the TGN^13,14^. This supports a model in which the two adaptors act independently, each regulating opposing transport routes^14^. Conversly, we found around 70% of all GGA domains to also harbour AP-1 (Fig. 5B), suggesting a functional connection of GGAs to AP-1. In addition, we observed dynamic and transient association and dissociation of both proteins from and to shared domains (Fig. 3D-E), suggesting their transient interaction may have a regulatory function. Intriguingly, in our proximome analysis (Fig. 6) we detected AP-1 specific cargoes exclusively in the AP-1, but not in any GGA TurboID experiment, suggesting that those proteins localize to the domains only in the absence of GGAs. Interestingly, the colocalization of GGAs with AP-1 increases on recycling compartments in comparison to secretory compartments (Fig. 4D). This raises the possibility that cargo association with AP-1 is regulated by GGAs, and that retrograde transport mediated by AP-1 occurs only when GGAs are absent from these domains. We speculate that GGAs mediate AP-1-independent anterograde transport from the TGN. In contrast, GGAs may associate with AP-1 on peripheral recycling compartments to modulate cargo retrieval, as reflected by their increased colocalization on transferrin-positive structures. Which molecular cues dictate association of AP-1 and GGAs and how GGAs would control retrograde transport via AP-1 remain elusive. Such a model would explain how competitive binding of both adaptors could orchestrate transport in opposing directions and integrates our results seamlessly into the existing literature^14,18^.

We found p56 and rabaptin-5 specifically enriched in the GGA proximome dataset in comparison to the AP-1 proximome (Fig. 6), indicating they may be required for GGA specific transport. Depletion of p56 led to a decreased motility of GGA carriers and an delay in processing of GGA cargo cathepsin D^30^, suggesting an involvement in GGA-mediated cargo export from the TGN. For rabaptin-5, it was shown that its binding to GGAs decreases GGAs capability to bind clathrin^41^. Rabaptin-5 acts in a complex with rabex5, the guanine-exchange factor of Rab5, and thus binding to rabaptin-5 could couple uncoating of GGA-carriers to fusion with a Rab5-positive target membrane. Whether p56 or rabaptin-5 regulate and possibly hinder the GGA-AP-1 interaction is not known.

Moreover, we found a smaller population of GGA domains devoid of clathrin (Fig. 5), suggesting that GGAs can associate and cluster to form domains without clathrin as a linker between molecules. However, we cannot conclude whether clathrin recruitment is required for GGA-mediated cargo transport. In addition, we observed clathrin-associated GGA domains with and without AP-1 (Fig. 5) which indicates that AP-1 is not required for clathrin recruitment to GGA sorting domains. This observation matches electron microscopy data from Drosophila cells where immune-gold labelling of AP-1 and GGA showed roughly 50% of the GGA vesicles also positive for AP-1^46^. These data support the notion that GGAs function both in an AP-1–dependent and an AP-1–independent manner, a mechanism conserved across species.

We found all three mammalian GGAs colocalized at TGN and in the cellular periphery (Fig. 1D). This finding is consistent with reports from others showing colocalization of GGAs^30,32^. We were curious whether simultaneous visualization of two endogenously-tagged GGAs would allow a more detailed understanding into potentially differences in regulation, dynamics or localization of different GGAs. However, we observed no differences in localization (Fig. 2), leaving it unclear to what extent the different GGAs are functionally different. Single GGA deletions did not affect localization of the other GGAs^30^, suggesting that they are recruited independently. While knock-out data from mice suggest at least for GGA2 a non-redundant function^9^, it is currently not clear which functionality this would be. Furthermore, it has been shown that depletion of GGA3 but not GGA1 or GGA2 defects β1-integrin trafficking^47^, a cargo that we find specifically enriched in AP-1 sorting domains (Fig. 6). Whether this observation is connected to a specific AP-1/GGA3 interaction or rather is caused by trafficking defect due to the absence of GGA3 remains unclear.

In summary, here we present new insights on how GGAs with AP-1 cooperate to regulate cargo selection and transport within human cells. Dynamic association and dissociation of adaptors domains suggests that interaction might have a regulatory role. The absence of AP-1–specific cargoes from shared domains led us to speculate that GGAs might regulate AP-1–mediated retrograde transport. We provide evidence shedding new light on current models of GGA- and AP-1–mediated transport^8,14^ and lay the groundwork for future studies focusing on the molecular interaction between these adaptors.

## Material and Methods

### Mammalian cell culture

CCL-2 HeLa cells (cat. no. 93021013 ECACC General Collection) were grown in a humidified incubator at 37 °C with 5% CO_2_ in Dulbecco’s Modified Eagle Medium (DMEM; Gibco) supplemented with 10% fetal bovine serum (Corning), 100 U l^−1^ penicillin and 0.1 g l^−1^ streptomycin (Fisher Scientific).

### Generation of CRISPR knock-in cell lines

### Cloning of guide and homology repair plasmids

### Guide plasmids

Guide RNAs were designed using the online tool Benchling (https://www.benchling.com) and cloned into the SpCas9 pX330 plasmid (Addgene plasmid #42230)^48^ by annealing oligonucleotides and ligation into the vector which was linearized with BbsI. A detailed list of guide RNA sequences is provided in Supplementary table 4.

### Cloning of HR plasmids

To avoid re-cutting from the Cas9, the protospacer-adjacent motif site was mutated in the HR plasmids. General design of HR plasmids and origin of the used sequences is described in^22,49^.

The following HR plasmid designs previously described: LoxP-G418^R^-LoxP-Halo-CLCa^EN 28^, AP1µA^EN^-SNAP-V5-PolyA-Puromycin^R 22^, AP1µA^EN^-Halo-ALFA-PolyA-G418^R 22^, ARF1^EN^-Halo-PolyA-G418 ^R 28^, ARF1^EN^-eGFP^22^.

### HR plasmids GGA1

The HR plasmid was designed with ∼1-kb homology arms and was synthesized by Twist Bioscience. A glycine–serine (GS) linker and a SpeI and EcoRI site were added between the two homology arms for insertion of tags and resistance cassette. To generate the GGA1^EN^-Halo-ALFA-PolyA-G418^R^ HR plasmid the entire insert (Halo-ALFA-PolyA-G418^R^) was amplified from the an existing AP1μA^EN^-Halo-ALFA-PolyA-G418^R^ HR plasmid^22^ using a SpeI Halo sense and G418 EcoRI antisense primer. The fragment was inserted into the ordered GGA1 HR plasmid, linearized with SpeI and EcoRI. To generate the GGA1^EN^-mStayGold-PolyA-Hygromycin^R^ HR plasmid the entire insert (mStayGold-PolyA-Hygromycin^R^) was amplified from the an existing AP1μA^EN^-mStayGold-PolyA-Hygromycin^R^ HR plasmid^22^ using a SpeI mStayGold sense and Hygromycin EcoRI antisense primer. The fragment was inserted into the ordered GGA1 HR plasmid, linearized with SpeI and EcoRI. To generate the GGA1^EN^-TurboID-V5-PolyA-G418^R^ HR plasmid the entire insert (TurboID-V5-PolyA-G418^R^) was amplified from the an existing AP1μA^EN^-TurboID-V5-PolyA-G418^R^ HR plasmid^35^ using a SpeI TurboID sense and G418 EcoRI antisense primer. The fragment was inserted into the ordered GGA1 HR plasmid, linearized with SpeI and EcoRI. To generate the GGA1^EN^-eGFP-PolyA-Puromycin^R^ HR plasmid, first, an eGFP-fragment was created by amplification of eGFP from a pEGFP-C1 plasmid using a BamHI GFP sense and a NheI antisense primer. This fragment was inserted into AP1μA^EN^-SNAP-V5-PolyA-Puro^R^ HR plasmid^22^ linearized with BamHI and EcoRI. In a second step the entire insert (eGFP-PolyA-Puromycin^R^) was amplified from the newly generated AP1μA^EN^-eGFP-PolyA-Puro^R^ HR plasmid using a SpeI eGFP sense and Puromycin EcoRI antisense primer. The fragment was inserted into the ordered GGA1 HR plasmid, linearized with SpeI and EcoRI. The extra step has become necessary as SpeI and NheI create identical restriction overhangs.

### HR plasmids GGA2

The HR plasmid was designed according to the design of the HR plasmid for GGA1 and was synthesized by Thermo Fisher. To generate the GGA2^EN^-Halo-ALFA-PolyA-G418^R^ HR plasmid the entire insert (SNAP-V5-PolyA-Puro^R^) was excised from the GGA1^EN^-Halo-ALFA-PolyA-G418^R^ HR plasmid using the SpeI and the EcoRI site and inserted into the ordered GGA2 plasmid, linearized with SpeI and EcoRI. Similarly, the GGA2^EN^-mStayGold-PolyA-Hygromycin^R^ HR plasmid, GGA2^EN^-TurboID-V5-PolyA-G418^R^ HR plasmid and the GGA2^EN^-eGFP-PolyA-Puromycin^R^ HR plasmid were created based of the existing GGA1 HR plasmids.

### HR plasmids GGA3

The HR plasmid was designed according to the design of the HR plasmid for GGA1 and was synthesized by Twist Bioscience. To generate the GGA2^EN^-Halo-ALFA-PolyA-G418^R^ HR plasmid the entire insert (SNAP-V5-PolyA-Puro^R^) was excised from the GGA1^EN^-Halo-ALFA-PolyA-G418^R^ HR plasmid using the SpeI and the EcoRI site and inserted into the ordered GGA2 plasmid, linearized with SpeI and EcoRI. Similarly, the GGA2^EN^-mStayGold-PolyA-Hygromycin^R^ HR plasmid, GGA2^EN^-TurboID-V5-PolyA-G418^R^ HR plasmid and the GGA2^EN^-eGFP-PolyA-Puromycin^R^ HR plasmid were created based of the existing GGA1 HR plasmids.

### Generation of CRISPR KI cell lines

CRISPR KI cell lines were created as described in ^49^. Briefly, HeLa cells at 70–80% confluency were transiently transfected with both guide and HR plasmids using FuGENE HD (Promega) according to the supplier’s protocol. Depending on the KI, G418, puromycin or hygromycin were added to the cells 3 days after transfection at a concentration of 1.5 mg ml^−1^ (G418) or µg ml^−1^ (puromycin) or 0.4 mg ml^−1^ (hygromycin) and the medium was exchanged every 2–days until selection was complete. KI cell lines were validated with western blotting. An overview of plasmids, base cell lines and selection method used for creation of new KI cell lines is given in Supplementary Table 5.

### Labelling for live-cell imaging

For all live-cell imaging experiments, cells were seeded on a glass-bottom dish (3.5 cm, no. 1.5; Cellvis) coated with fibronectin (Sigma) one day prior to the experiment. Labelling with HaloTag and SNAP-tag substrates (substrate concentration: 1 µM) was carried out for 1 h at 37 °C in culture medium. All live-cell imaging dyes used in the study are indicated in the figure legends and were provided from the Lavis lab^50,51^. After the staining, cells were washed in growth medium at 37 °C for at least 1 h. Live-cell imaging was performed in live-cell imaging solution (FluoroBrite DMEM (Gibco) supplemented with 10% FBS, 20 mM HEPES (Gibco) and 1× GlutaMAX (Gibco)).

### Imaging and image processing

Line-scanning confocal microscopy data were collected on a commercial expert line Abberior STED microscope equipped with 485 nm, 561 nm and 640 nm excitation lasers using the Imspector software from Abberior Instruments (version 16.3). For confocal imaging, probes that were detected in the 498 to 551 nm and the 650 to 756 nm detection windows were recorded simultaneously. If required for quantitative analysis, the laser power was kept constant between images. The pixel size was set to 60 nm and live-cell imaging was performed at 37 °C.

Spinning-disk confocal imaging was carried out using a CSU-W1 SoRa spinning disk (Nikon) with NIS-Elements software (version 4.50). A dual camera system for simultaneous dual-colour detection was used. Imaging was performed with a ×60 Plan Apo oil objective (NA = 1.4) and the 488 nm, 561 nm and 636 nm laser lines were used for excitation. The eGFP and far-red (JFX_650_) signals were detected simultaneously, and the orange channel (JF_552_) was detected separately. A quad-bandpass filter was used with relevant centre wavelengths of 521 nm, 607 nm and 700 nm and full-width half maximums of 21 nm, 34 nm and 45 nm, respectively.

To reduce noise, confocal images were background subtracted and Gaussian blurred (1 pixel s.d.) using Fiji^52^.

### Trafficking assays

For the RUSH assay (Fig. 4A) a SBP-M6PR(RUSH)-SNAP plasmid was transiently transfected into HeLa cells. 1 day after transfection, cells were labelled with BG-JF_552_ and BG-JFX_650_. The biotin stock solution (c = 585 mM in dimethylsulfoxide) was diluted to a final concentration of 500 µM in live-cell imaging solution and then added to the cells. Live-cell confocal imaging was started 15 min after biotin addition to investigate post-Golgi trafficking of the cargo and cells were imaged every 2 min.

For the Tfn assays (Fig. 4B-D), live-cell labelled HeLa cells were put on ice for 10–15 min before incubated with Tfn-AF555 (25 µM, Thermo Fisher) in live-cell imaging solution at 37 °C for 2 min. Cells were washed once with live-cell imaging solution before live-cell imaging with holo-Tfn (1 mM, Thermo Fisher) was added to prevent re-endocytosis of labelled Tfn. Cells were imaged 5-10 min after addition of Tfn.

### Image quantification

To quantify GGA intensity at Golgi and peripheral ARF1 compartments (Fig. 1C), confocal images of ARF1^EN^-Halo/GGA2 ^EN^-eGFP HeLa cells were analysed. Each cell was imaged at the PM plane and the Golgi plane. In each cell the fluorescence intensity of ten GGA puncta which were associated with ARF1 compartments were measured and averaged, the same was done for GGA puncta at the Golgi. The signals were then normalized to the averaged Golgi-GGA intensity.

Colocalization of GGA2 with AP-1 on Tfn-positive compartments (Fig. 4D) was done by calculating the Manders correlation coefficient. Confocal images of GGA2^EN^-mStayGold/AP1µA^EN^-SNAP HeLa cells 5-10 min after Tfn-AF555 addition (as described for Fig. 4B) were Gassian blurred and background substracted. For each channel a threshold was manually chosen and the Tfn signal was converted into a mask. This mask was used to determine the Manders correlation coefficient of GGA2 and AP-1 using the Coloc 2 plugin of Fiji. An inverted mask of the Tfn signal was used to determine the colocalization of GGA2 and AP-1 which do not localize on Tfn compartments. For overall colocalization analysis no ROI was chosen. Statistical analysis was performed in GraphPad Prism (GraphPad Software; https://www.graphpad.com).

To estimate the association of GGAs with clathrin and AP-1 (Fig. 5B), all GGA punctae were manually counted in 10 GGA2^EN^-mStayGold/AP1µA^EN^-SNAP/Halo-CLCa^EN^ HeLa cells imaged with scanning confocal microscopy from three independent experiments and categorized into three categories depending on their colocalization with either clathrin without AP-1, clathrin with AP-1 and or neither of them.

### Preparation of MS samples

MS samples were prepared as described previously^35^. Briefly, for each of four replicates, cells of the different KI cell lines (AP1µA^EN^–TurboID^35^, GGA1^EN^–TurboID, GGA2^EN^–TurboID and GGA3^EN^–TurboID) were seeded into two 15 cm cell culture dishes (1.5 million cells per dish). As control, HeLa WT cells were used and seeded in the same way. At 24 h after seeding, the medium was replaced with medium supplemented with 50 µM biotin for another 24 h. All samples were washed five times with ice-cold PBS and then detached in 4 ml PBS and collected in a falcon tube. Cells were pelleted by centrifugation at 300 g at 4°C for 3 min. The supernatant was removed and the pellet resuspended in 4 ml RIPA lysis buffer (50 mM Tris-HCl pH 7.4, 150 mM NaCl, 0.1% SDS, 0.5% sodium deoxycholate and 1% Triton X-100) supplemented with 1× protease inhibitors (Roche). To ensure sufficient cell lysis and full disruption of membranes, all pellets were lysed for 30 min on ice and cell membranes were disrupted by passing the cells 10 times (five strokes) through a 24 G needle. The cell lysates then were centrifuged in microcentrifuge tubes at 13,000 g at 4°C for 10 min. Next, clarified lysates were mixed with 150 µl of streptavidin magnetic beads which were equilibrated twice with RIPA lysis buffer. The samples were incubated with the magnetic beads rotating at 4°C overnight. On the next day, the beads were washed twice with RIPA lysis buffer, once with 1 M KCl followed by quick washes with 0.1 M Na_2_CO_3_ and 2 M urea in 10 mM Tris-HCl (pH 8.0). The beads were then washed twice in RIPA lysis buffer and transferred into fresh microcentrifuge tubes. Subsequently, they were washed once with 50 mM Tris-HCl (pH 7.4) and twice with 2 M urea in 50 mM Tris-HCl (pH 7.4). The wash buffer was removed, and the beads were resuspended in 80 µl of 2 M urea in 50 mM Tris-HCl (pH 7.4) with 1 mM DTT and 0.4 µg trypsin and incubated for 1 h shaking at 1000 rpm. The supernatant was transferred into fresh microcentrifuge tubes and the beads were washed twice with 60 µl of 2 M urea in 50 mM Tris-HCl (pH 7.4). The washes were combined with the supernatant, DTT was added to a final concentration of 4 mM and the samples were incubated for 30 min at 25°C shaking at 1000 rpm. Iodoacetamide was added to final concentration of 10 mM and the samples were incubated in the dark at 25°C for 45 min while shaking at 1000 rpm. Finally, another 0.5 µg of trypsin were added and the digestion was proceeded overnight at 25°C shaking at 700 r.p.m. On the next day, the digestion was stopped by acidification with formic acid to a final concentration of 1% (v/v). The peptides were prepared for MS using SDB-stage tips as described in ^35^.

### Nano LC-MS and data analysis

Peptides were reconstituted in 10 µl of 0.05% trifluoroacetic acid (TFA), 4% acetonitrile and 5 µl were analyzed by an Ultimate 3000 reversed-phase capillary nano liquid chromatography system connected to a Q Exactive HF mass spectrometer (Thermo Fisher Scientific). Samples were injected and concentrated on a trap column (PepMap100 C18, 3 µm, 100 Å, 75 µm i.d.×2 cm, Thermo Fisher Scientific) equilibrated with 0.05% TFA in water. After switching the trap column inline, LC separations were performed on a capillary column (Acclaim PepMap100 C18, 2 µm, 100 Å, 75 µm i.d.×25 cm, Thermo Fisher Scientific) at an eluent flow rate of 300 nl/min. Mobile phase A contained 0.1% formic acid in water, and mobile phase B contained 0.1% formic acid in 80% acetonitrile and 20% water. The column was pre-equilibrated with 5% mobile phase B followed by an increase of 5–44% mobile phase B over 100 min. Mass spectra were acquired in a data-dependent mode utilizing a single MS survey scan (*m*/*z* 300–1650) with a resolution of 60,000, and MS/MS scans of the 15 most intense precursor ions with a resolution of 15,000. The dynamic exclusion time was set to 20 s and automatic gain control was set to 3×10^6^ and 1×10^5^ for MS and MS/MS scans, respectively. The mass spectrometry proteomics data were deposited to the ProteomeXchange Consortium via the PRIDE partner repository^53^ with the dataset identifier PXD074768.

MS and MS/MS raw data were analyzed using the MaxQuant software package (version 2.0.2.0). Data were searched against the human reference proteome downloaded from Uniprot (82,678 proteins, taxonomy 9606, last modified October, 2023) using the default parameters except for enabling the options ‘label-free quantification (LFQ)’ and ‘match between runs’. Filtering and statistical analysis was carried out using the software Perseus version 1.6.14^54^.

Only proteins which were identified with LFQ intensity values in in at least three out of four replicates were used for downstream analysis. Missing values were replaced from normal distribution (imputation) using the default settings in Perseus (width 0.3, down shift 1.8). Mean log_2_ fold protein LFQ intensity differences between experimental groups calculated in Perseus using unpaired two-tailed Student’s *t*-tests with a permutation-based false discovery rate (FDR) of 0.05 to generate the adjusted *P*-values (q-values).

### Volcano plots

Volcano plots were created by plotting the −log_10_ *P*-value against the mean log_2_ fold protein LFQ intensity differences. The volcano plots include all proteins that were detected for AP-1 and the indicated GGA.

### Proximome analysis

For proximome analysis of GGAs and AP-1 (Supplementary tables 1-3), only proteins that were significantly enriched (*P*<0.05 and log_2_ fold change >1) against the WT control sample were considered. Proteins that were additionally enriched in GGA samples vs AP-1 samples and vice versa were then screened with UniProt to identify proteins that were known to localize to either the Golgi or endosomal membranes, act in or regulate cellular transport, or are transmembrane proteins and thus could be cargoes of the studied adaptors. The results of this analysis are shown in supplementary table 1 (Proteins enriched for AP-1) and supplementary table 3 (Proteins enriched for the different GGAs). Proteins that were significantly enriched in GGA and AP-1 samples compared to the WT control were considered common interactors and are summarized in supplementary table 2. These tables are not mutually exclusive as proteins that are listed in supplementary table 2 can also be part of supplementary table 1 or 3.

### Statistics and reproducibility

GraphPad Prism 8.0.2 was used to generate all graphs. All schematics were generated with Affinity Designer 2. Microscopy images and western blots are shown as representative images. Micrographs in Figs. 1A-B, 2A-C, 3A-E, 4A-C, and 5A are representatives of 3 independent experiments.

## Acknowledgements

This project was supported by the Deutsche Forschungsgemeinschaft (DFG, German Research Foundation) grant SFB958 (project A25 to F.B.) and DFG – Project Number 278001972 – TRR 186. For mass spectrometry, we would like to acknowledge the assistance of the Core Facility BioSupraMol supported by the DFG. We thank Scottie Robinson for feedback on the manuscript. We thank L. Lavis (Janelia Research Campus) for providing all live-cell dyes we used in this study. We thank S. Donat and J. Schmoranzer of the AMBIO imaging centre (Charité Berlin) for the technical assistance for spinning-disk experiments.

## Author Contributions

A.S. and F.B. conceived the project. A.S., A.K., A.D., L.S.B., J.M. and F.B. designed and performed experiments. A.S., A.K. and A.D. performed image analysis. C.F. and B.K. assisted with MS-experiments. A.S., A.K., A.D., and J.M. generated plasmids and knock in cell lines. A.S. and F.B. wrote the manuscript with input from all authors.

---

**Supplementary Figure 1:**
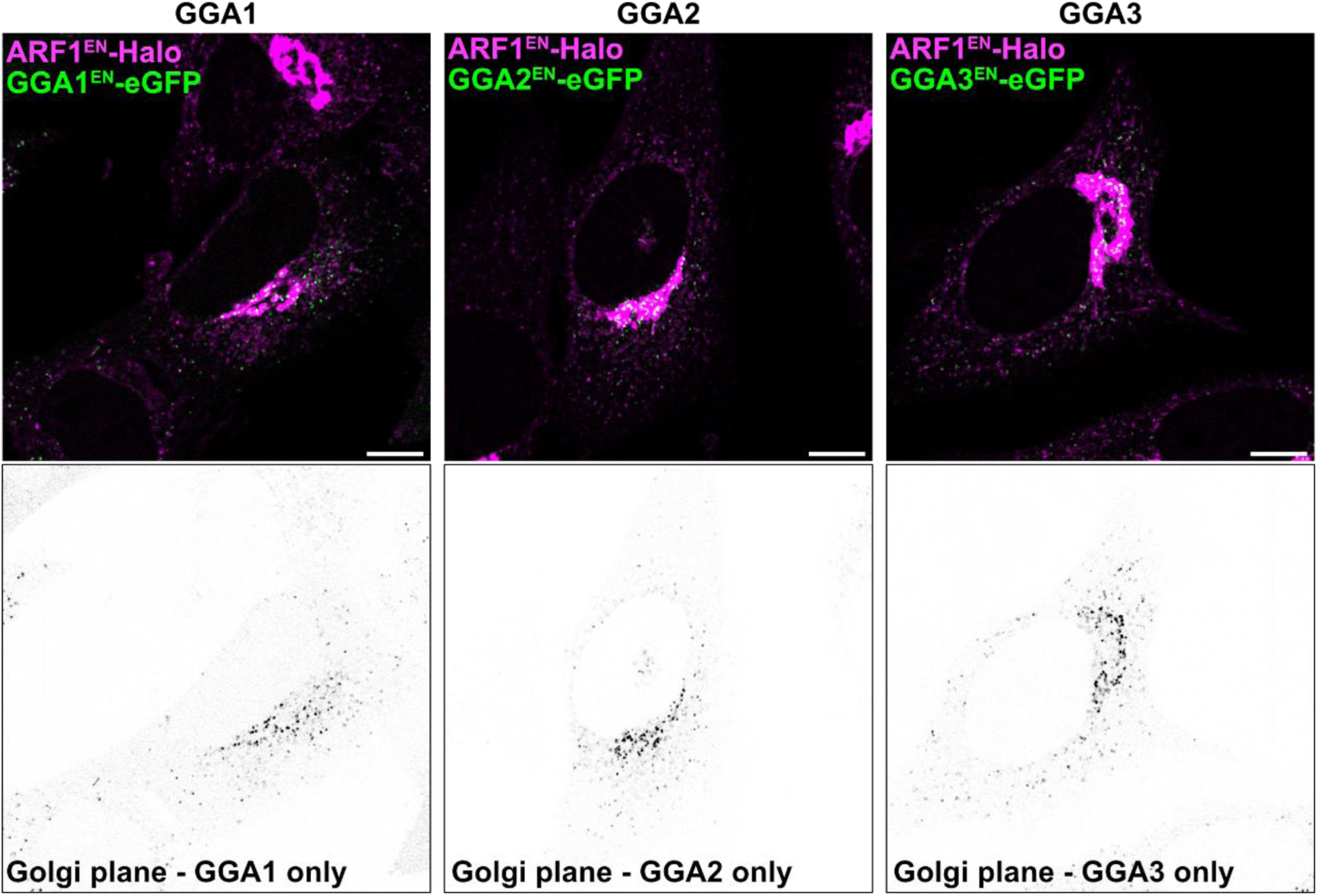
GGAs localize to Golgi membranes. Live-cell confocal imaging of GGA1^EN^-eGFP/ARF1^EN^-Halo, GGA2^EN^-eGFP/ARF1^EN^-Halo and GGA3^EN^-eGFP/ARF1^EN^-Halo HeLa cells labeled with CA-JFX_650_ shows GGAs association to the Golgi and TGN when focussing the plane that resolves the Golgi best. Single-color images of the different GGAs highlight their localization in the Golgi area. Scale bars: 10 μm.

**Supplementary Figure 2:**
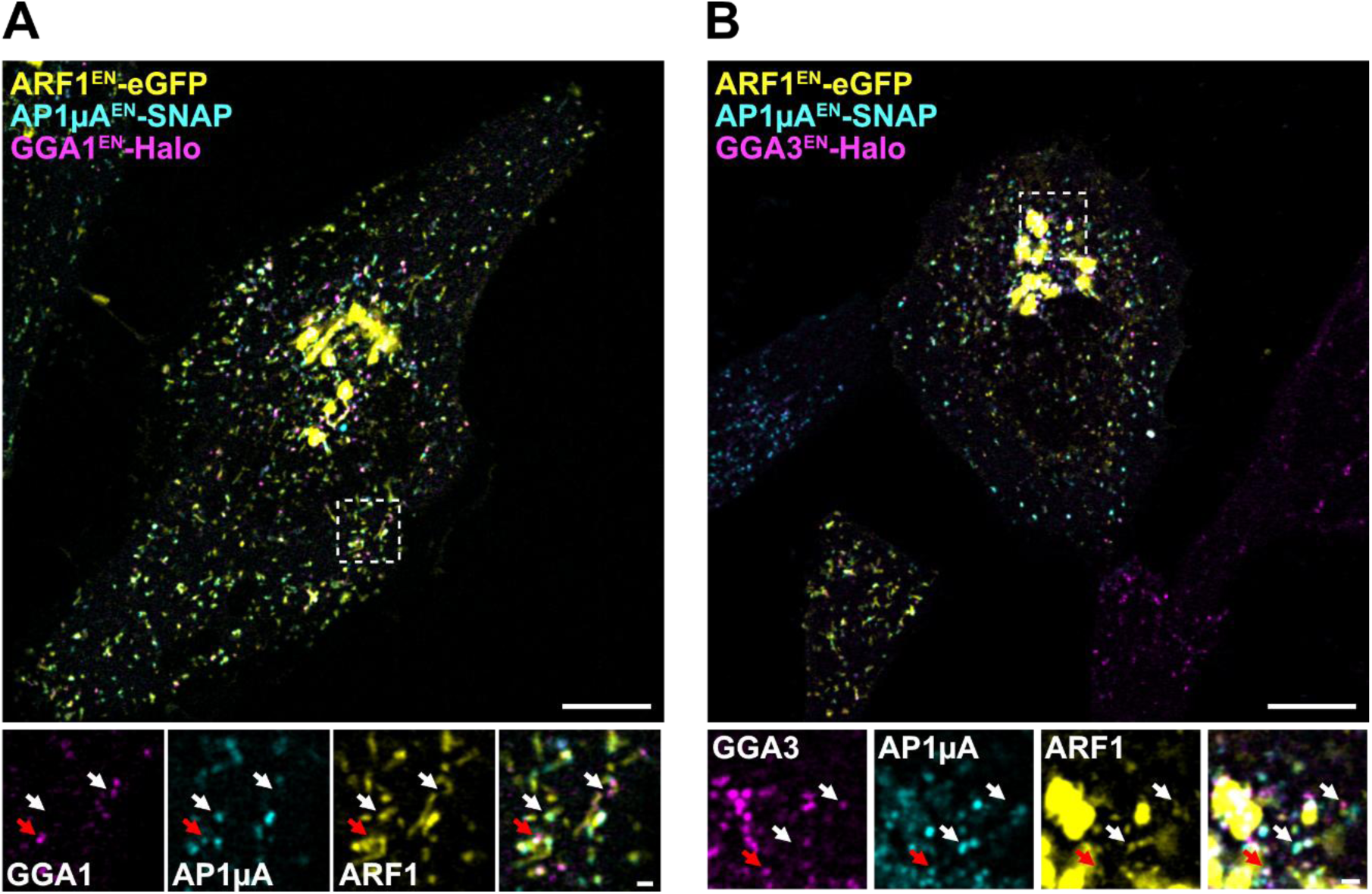
Transient association and dissociation of AP-1 and GGAs from shared membrane domains. (**A**) Live-cell confocal imaging of ARF1^EN^-eGFP/AP1µA^EN^-SNAP/GGA1^EN^-Halo HeLa cells labeled with BG-JF_552_ and CA-JFX_650_. (**B**) Live-cell confocal imaging of ARF1^EN^-eGFP/AP1µA^EN^-SNAP/GGA1^EN^-Halo HeLa cells labeled with BG-JF_552_ and CA-JFX_650_. Magnifications highlight domains at the TGN (in B) and at peripheral ARF1 compartments (in A) where only one of the two adaptors AP-1 or GGA2 (highlighted with white arrows) is present. Also, domains were both adaptors were present were found (highlighted with red arrows). Scale bars: 10 μm (overviews) and 1 µm (crops).

